# Big DNA as a tool to dissect an age-related macular degeneration-associated haplotype

**DOI:** 10.1101/461251

**Authors:** Jon M Laurent, Xin Fu, Sergei German, Matthew T Maurano, Kang Zhang, Jef D Boeke

## Abstract

Age-related Macular Degeneration (AMD) is a leading cause of blindness in the developed world, especially in aging populations, and is therefore an important target for new therapeutic development. Recently, there have been several studies demonstrating strong associations between AMD and sites of heritable genetic variation at multiple loci, including a highly significant association at 10q26. The 10q26 risk region contains two genes, *HTRA1* and *ARMS2*, both of which have been separately implicated as causative for the disease, as well as dozens of sites of non-coding variation. To date, no studies have successfully pinpointed which of these variant sites are functional in AMD, nor definitively identified which genes in the region are targets of such regulatory variation. In order to efficiently decipher which sites are functional in AMD phenotypes, we describe a general framework for combinatorial assembly of large ‘synthetic haplotypes’ along with delivery to relevant disease cell types for downstream functional analysis. We demonstrate the successful and highly efficient assembly of a first-draft 119kb wild-type ‘assemblon’ covering the *HTRA1/ARMS2* risk region. We further propose the parallelized assembly of a library of combinatorial variant synthetic haplotypes covering the region, delivery and analysis of which will identify functional sites and their effects, leading to an improved understanding of AMD development. We anticipate that the methodology proposed here is highly generalizable towards the difficult problem of identifying truly functional variants from those discovered via GWAS or other genetic association studies.

## Introduction

### Age-related macular degeneration is a complex genetic disease characterized by common haplotype associations

Age-related macular degeneration (AMD) is considered to be the leading cause of vision loss in the elderly of developed countries (1-3). As the populations of these countries continue to age, AMD will increase in prevalence, and its already significant contribution to health care costs will continue to grow. The disease is characterized by abnormalities in the retinal pigmented epithelium (RPE), which underlies the photoreceptor cells of the retina (**Fig. 1**) (4). AMD primarily affects the macula, a region of the central retina associated with the highest acuity in the visual field. While the etiology of AMD is complex and unclear, the disease has a well described progression from early, to late, and on to advanced forms. The advanced forms of the disease are the primary cause of vision impairment due to AMD, with 90% of patients exhibiting severe central vision loss having progressed to advanced stages of disease (5).

**Figure 1.**
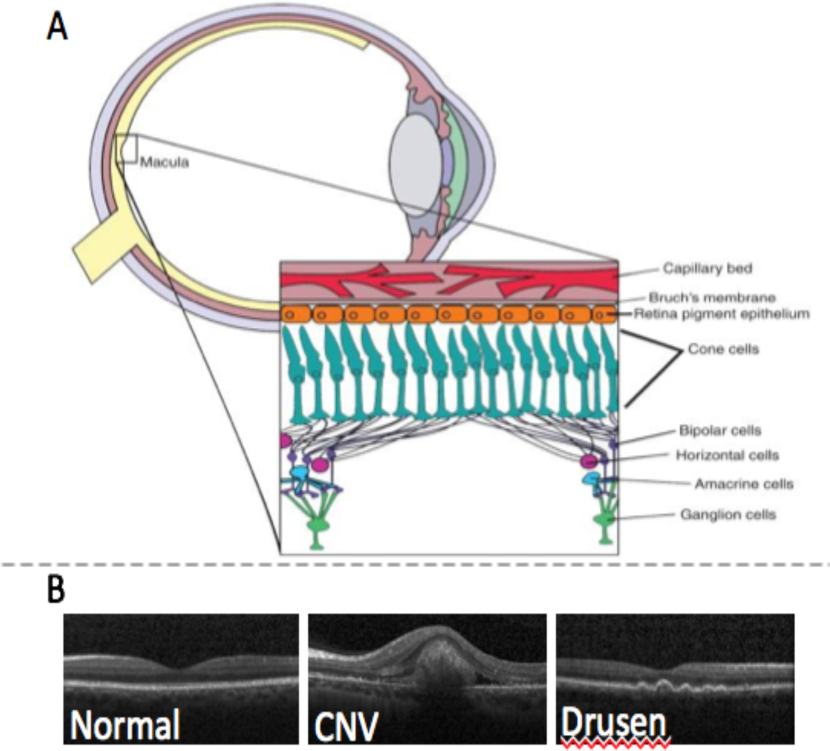
Pathology of AMD. **(A)** Schematic of eye showing tissues affected by macular degeneration. **(B)** Optical coherence tomography images of a normal (left), AMD retina showing choroidal neovascularization (center), and AMD retina showing drusen deposits (right).

Despite the well-known progression of disease stages, AMD is not curable, and current therapeutic interventions have had limited effectiveness, owing largely to its complex genetics and incomplete knowledge of the disease’s underlying mechanisms. A combination of high-dose zinc along with vitamins C and E and β-carotene is a common treatment recommended for newly diagnosed patients (6). The treatment has no effect on developing AMD but may have some benefit for slowing progression of AMD once it has started. Vascular endothelial growth factor (VEGF) is a major factor associated with new blood vessel growth and is thus expressed in the neovascular membranes of wet AMD. Anti-VEGF therapies have therefore been used as some of the most effective treatments for CNV, with up to 90% of patients seeing stabilized vision and nearly a third reporting significantly improved vision in some clinical trials (7, 8).

Both environmental and genetic risk factors have been implicated in the development and progression of AMD. Smoking is the strongest modifiable environmental contributor to AMD, and has been shown to increase the overall risk of AMD diagnosis by two-to three-times (9). However, environmental risk factors are likely only modifiers of existing genetic predisposition to AMD, and indeed we and others have performed genome-wide association studies (GWAS) to demonstrate that development and progression of AMD is strongly tied to a number of loci, such as the complement pathway genes complement factor H (CFH), C3, CFB, and CFI (10-14). Another highly significant AMD-associated region has been identified and corroborated extensively (15-18), and lies on chromosome 10 at 10q26. Unlike the monogenic complement loci associated with AMD, the 10q26 region includes parts of three different genes, each of which could be associated with AMD. The protein encoded by one of the genes, *HTRA1*, is found in drusen deposits, and is therefore an attractive causal agent, though conflicting results have implicated both it and the adjacent *ARMS2* as being affected by the AMD-associated risk haplotypes in the region.

### The HTRA1/ARMS2 locus

The significant AMD susceptibility locus at 10q26 contains three candidate target genes, *PLEKHA1, ARMS2* and *HTRA1* **(Fig. 2)**. Many sequence variants have been found in the region, contained within a number of risk and protective haplotypes. The Zhang lab recently resequenced a ∼100kb region surrounding *HTRA1*, a secreted serine protease, and *ARMS2*, a gene of unknown function, in a group of AMD patients and controls, which resulted in identification of 67 variants in the region. A follow-up assocation study with a 200 case, 200 control cohort refined the likely primary risk haplotype to a 21.5kb region including parts of both *HTRA1* and *ARMS2* and encompassing 22 variants (19). Thorough characterization has suggested that the primary risk and protective haplotypes can be distinguished by the identity of alleles present at four of these variants. The current literature presents contradictory evidence for which of these genes is the target of regulatory variants in the region. Early results suggested variants in the promoter region of *HTRA1* led to increased expression and were associated with an increased incidence of AMD (17, 18). Subsequent studies were unable to corroborate these results, and *ARMS2* was again implicated as potentially causative (20). More detailed investigation identified several new polymorphisms associated with a decrease in ARMS2 transcript levels and increased likelihood of developing AMD (21). Still further analysis has posited that a combination of the two effects (loss of *ARMS2* transcript and increased *HTRA1* expression) is necessary (19). Thus, in order to identify the functional sites and targets of the risk haplotype, it is necessary to exhaustively assay all variants within the region, alone and in combination, and characterize their downstream effects.

**Figure 2.**
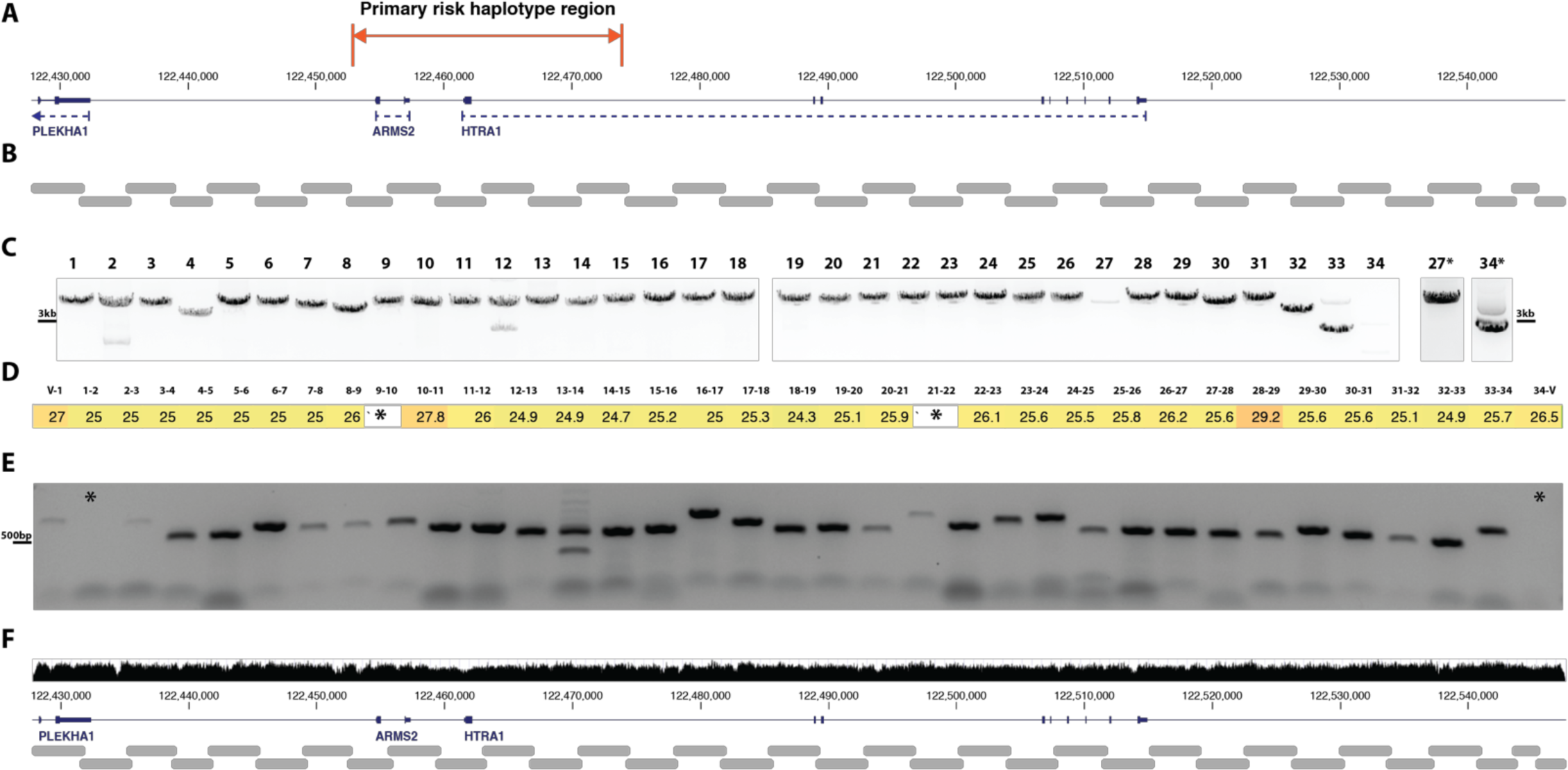
Assembly scheme for initial HTRA1/ARMS2 assemblon. (**A)** The genome locus surrounding HTRA1 and ARMS2 that was assembled here. **(B)** Approximate size and position of the assembly segments designed by our automated platform. **(C)** PCR of assembly segments from a BAC clone covering the region. Segments 27 and 34 were amplified separately with alternative conditions, shown with *. **(D)** Example output from the automated RT-PCR junction PCR strategy showing crossing-point data and **(E)** manual junction PCR strategy for the same assemblon clone. * indicates junctions that did not amplify in the respective assay, but were ultimately present. **(F)** Sequence coverage track of the same assemblon clone in D and E obtained by WGS of the yeast strain.

### Functional dissection of genomic variation with big DNA

Nuclease-based genome engineering methods like CRISPR-Cas9 have come to the fore recently, but are less than ideal for high-throughput studies of multiple loci or those that require large genomic alterations, as they are capable of only one to a few edits at a time, as well as prone to off-targeting and undesirable repair outcomes. Testing many variants, especially in combination, is thus labor-and cost-prohibitive. Further, in diploid genomes, such top-down editing methods require extensive screening of clones in order to identify those with proper mutational phasing (scaling as 2^n^ clones for n mutations). Our bottom-up approach, in contrast, allows pairs or higher order combinations of variants of interest to be combined in cis in a single step, rather than randomly across homologs. In general, such methods for large-scale ‘writing’ and wholesale replacement of large genomic sequences will enable rapid high-throughput assessment of many sequence variants either in isolation or combined in cis and importantly, many permutations can be built and assayed efficiently in parallel. We have been pioneering such technologies for the past several years, most notably in the design and synthesis of an entire eukaryotic genome with the Sc2.0 consortium (22). One such method for assembly of large DNA constructs is called “SWAP-In” and has been used to deliver segments of DNA ranging in length from 30,000 to 60,000 bp precisely to the desired location in the yeast genome by exploiting the endogenous homologous recombination (HR) mechanism in yeast. Moreover, this process could be iterated to produce synthetic yeast chromosomes of up to nearly 1 Mb (22-29). We are now applying these methods towards assembly of large mammalian loci and pathways called “assemblons” because they are assembled by HR in the yeast cell. These assemblons are built by a specific instantiation of our homologous recombination-based assembly method called extrachromosomal SWAP-In (eSWAP-In) that we recently described (30). Importantly, these assemblons can be designed such that they allow combinatorial assembly of one or multiple variant segments interspersed with wild-type sequence segments.

In recent work (30), we have shown that it is possible to amplify a series of overlapping DNAs spanning the human *HPRT1* locus, and using eSWAP-In, rapidly assemble a 101 kb intact human *HPRT1* region in three sequential eSWAP-In steps. The resulting plasmid (essentially a designer BAC) was recovered ino *E*. *coli* cells, and the large insert was analyzed by restriction digests and shown to have the correct structure. The resulting insert DNA, along with flanking plasmid sequences was integrated into the X chromosome of mouse embryoninc stem cells (ESCs), near the native murine Hprt1 locus. DNA sequencing established that the entire human locus was delivered intact at the expected position. Immunoblotting established that the human gene was expressed in the ESCs.

### Defining the impact of synthetic haplotypes

Like most disease-associated variants identified by GWAS, most of the variants reported in the AMD-associated region in 10q26 lie in non-coding regions, either intergenic or intronic to the genes located there. Thus, the functions of these variants are likely regulatory in nature, either cis-acting at the nearby genes or potentially trans-acting across the genome. We will therefore use gene expression measurements as the initial means for assessing functional impact of the AMD associated haplotypes. Gene expression effects will first be assayed by local measurements of nearby genes (i.e. *HTRA1, ARMS2*, and *PLEKHA1*), as these are the most likely targets of regulatory variants in the region, but can be expanded to include transcriptome-wide measurements. More detailed cellular effects can be investigated as well, such as chromatin accessibility or transcription factor occupancy.

We will tackle this problem by building an assemblon consisting of the entire *HTRA1/ARMS2* locus and a substantial swath of flanking DNA, encompassing nearly 120 kilobases. Once assembly is demonstrated, a library of variant assemblons can be rapidly produced using using the same strategy. The variant assemblon library will be designed to allow functional interrogation of all variants in the region, as well as combinatorial analysis of multiple variants, to identify possible epistatic interactions between them **(Fig 3)**. For example, suppose there are ten SNVs in linkage disequilibrium with the index risk SNV. It would be simple to prepare ten variants each bearing a single risk SNV, and 40 additional variants bearing all possible combinations of two SNVs. These can then be delivered to a suitable test cell, such as a retinal pigment epithelial (RPE) cell homozygous for the protective haplotype, replacing one of those “wild-type” alleles with a landing pad similar to that described by (30). This would allow the 50 engineered loci to replace one of the endogenous alleles in a single step. It would be important to “tag” the two alleles, e.g. with a fluorescent protein gene or a natural mRNA variant in order to distinguish the engineered alleles from the endogenous protective allele. Assuming the engineered allele is tagged, e.g. HTRA1-GFP and ARMS2-RFP, the readout of gene expression would be straightforward. Importantly, it is possible that this GWAS locus, and perhaps many others, is driven not by a single variant but by epistasic relationships between variants, and perhaps even expression effects on more than one gene. Indeed, in the case of *HTRA1/ARMS2*, multiple previous studies have implicated both genes separately as the target of AMD-associated regulatory variants, as well as suggesting multiple variants to be functional (20, 21, 31, 32).

**Figure 3.**
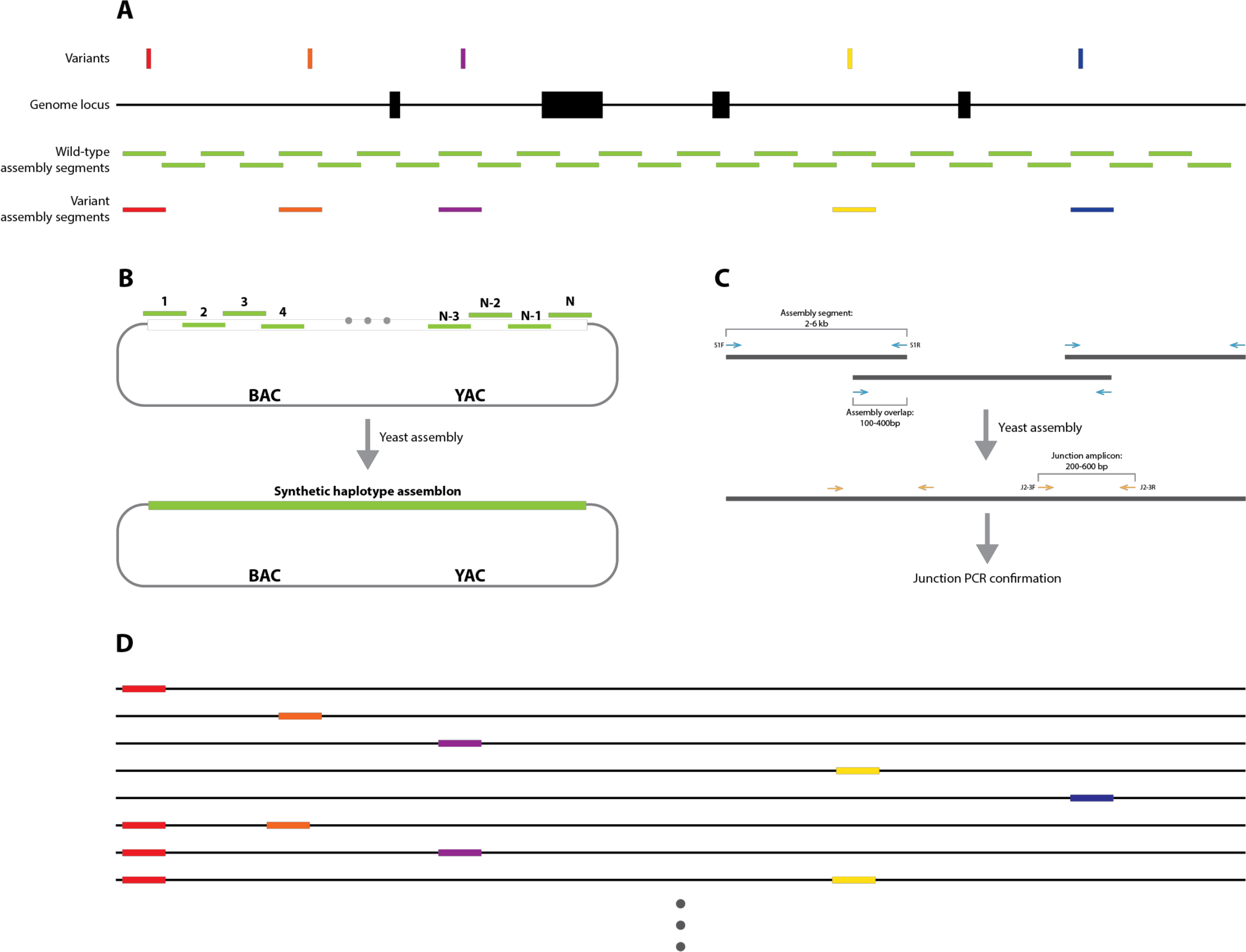
Generic scheme for interrogating disease-associated genetic variation with synthetic haplotypes. (**A)** A genome locus associated with some disease or phenotype, showing locations of associated variants, overlapping assembly segments, for wild-type and variant assembly. **(B)** Outline of the assembly method. Overlapping assembly segments are co-transformed into yeast along with a linear assembly vector, where they are efficiently assembled into a complete assemblon **(C)** Assembly segments are initially obtained as PCR amplicons or synthetic segments using ‘segment primers’. Correct assembly structure is first screened by PCR across the assembly junctions with ‘junction primers’ **(D)** A subset of possible variant synthetic haplotypes that would be present in the assembled variant library for the theoretical locus shown in **A**. These can be efficiently assembled and delivered in parallel, allowing high-throughput interrogation of which variant sites are functional.

## Results and Discussion

### Assembly of a ‘first-draft’ 119 kb HTRA1/ARMS2 locus

With the aim of demonstrating the feasibility of our approach, we have completed a ‘first-draft’ assembly of the wild-type *HTRA1/ARMS2* locus. We chose a ∼119kb region that encompasses both the *HTRA1* and *ARMS2* genes and the primary AMD risk haplotype identified previously (19) **(Fig. 2A**), as well as a significant portion of up-and downstream sequence. The region was broken up into 34 segments averaging ∼3.8 kilobases in length (2.1-4.1 kb), overlapping by an average of 370bp **(Fig. 2B-C)**. Segments were obtained by amplification from a commercially available BAC clone containing the entire region. PCR amplicons were purified and co-transformed into yeast along with a linearized ‘assembly vector’ containing sequences overlapping the first and last assembly segments **(Fig. 3B)**. Importantly, assembly of the *HTRA1/ARMS2* assemblon was carried out in a single step rather than iterative assembly, meaning construction of the variant assemblon library will be efficiently parallelized. We identified complete assemblies by amplifying across the assembly junctions in an automated RT-PCR workflow **(Fig. 2D-E)** (30). Several dozen clones with the correct structure by junction PCR were fully sequence confirmed by whole-genome sequencing of the yeast strain **(Fig. 2F)**.

The proposed combinatorial assembly of “synthetic haplotypes” will allow us to assess the regulatory impact of many variants within the risk region in isolation or combination, and thus in principle we will be able to narrow down and identify those variants likely to contribute to AMD development and progression. We can further characterize the functional impact of these sites by interrogating their effects on chromatin accessibility and transcription factor binding in relevant cell types.

### The bigger picture

GWAS studies have identified thousands of loci associated with thousands of diseases or other phenotypes (33). In the vast majority of cases (>90%), the risk SNVs (single nucleotide variants) fall outside of coding exons, and are thus likely phenotypically associated through some unknown regulatory function (34). Generally, these risk SNVs are in linkage disequilibrium (LD) with a substantial number of other local SNVs and other types of molecular variants such as indels, meaning they are typically inherited as part of a “haplotype block” tens to hundreds of kilobases in size. The correlation between variants in the associated region makes teasing apart causal variants difficult and unsatisying, as the set of risk-associated SNVs is rarely broken up by recombination events occuring during meiosis. Given the number of potentially causative variants correlated with typical GWAS index variants, not to mention possible epistatic effects, current ‘top-down’ genome editing techniques leave much to be desired in attempts to identify the truly functional sites in these haplotypes.

We thus propose that GWAS hits can be more efficiently dissected by use of synthetic haplotype ‘writing’ and subsequent analysis of engineered human cell lines, outlined above for AMD, as a general method summarized in **Fig. 3**. In many diseases, the cell types of interest are diffcult to access or difficult to transfect. In these cases, we can make use of induced pluripotent stem cells (iPSCs) as the engineering platform, and then induce differentation towards a relevant disease cell type. Extending this idea further would make use of mouse embyonic stem cells (ESCs). Mouse ESCs have one very special advantage in that they can be developed completely into animals, from which organismal phenotypes can be measured, or any cell type can be analyzed or harvested. We and others have shown that human DNA sequences larger than 100 kilobases can be delivered to mESCs and not only support expression (30) but development of viable animals. As shown for the alpha-globin locus by Doug Higgs and his colleagues (35), human genes are even expressed only in the relevant cell types, albeit at a slightly lower level than the corresponding mouse gene. Related work by Duncan Odom, in which an entire human chromosome 21 was transferred to mESCs (36), showed that not only was the human chromosome segregated and replicated normally in the animal, but the genes were expressed in an accurately tissue specific manner. These two remarkable studies demonstrate that all or most of the regulatory sequences necessary for proper gene expression, including tissue-and organ-specificity, are highly conserved between mouse and human in a functional sense, implying that the transacting factors that recognize them can effectively reach across substantial species boundaries with great specificity.

Once full risk and protective haplotypes are transferred to stem cells, and either differentiated into an appropriate cell type or converted to mice, the observed transcription differences between them should provide the information needed to develop a transcription-based assay for the variant synthetic haplotypes. With this tool in hand, the variant haplotypes can then be analyzed to fine-map the causative functional variant(s).

## Methods

### Assembly region definition

The region for assembly was chosen somewhat arbitrarily while fulfilling a specific set of criteria. Namely, it had to encompass the previously identified primary AMD risk haplotype as well as the larger surrounding region resequenced by the Zhang lab (19). It also contained the entirety of the *HTRA1* and *ARMS2* genes, and a substantial portion of upstream and downstream sequence in order to capture as much potential regulatory sequence as possible. The ends of the region were further chosen to not interfere with any possible regulatory sites demarkated by histone modification or DNaseI hypersensitivity peaks as shown in the UCSC genome browser.

### Primer design

Primers for segment amplification as well as assembly junction screening were designed using an in-house developed algorithm. The algorithm chooses assembly segment amplification primers in such a way that they will have minimal off-target binding in the genome of choice, while amplifying appropriatly sized segments for efficient assembly. Junction screening primers are chosen such that they span the assembly junction (i.e. will not produce a product unless two segments have fused), and produce an amplicon that can be efficiently amplified from a ‘dirty’ genomic DNA prep (generally <500bp).

### Assembly segment preparation

Most assembly segments were amplified from a commercially available BAC (clone RP11-467N7 from the Chori BACPAC resources center, https://bacpacresources.org), using KAPA HiFi HotStart DNA polymerase (KAPA Biosystems). 32/34 segments were successfully amplified with default KAPA cycling conditions, and an annealing temperature of 65°C and extension temperature of 68°C. Segment 27 was successfully amplified with a 65°C extension temperature. Amplicon 34 required use of Q5 polymerse (New England Biosciences) to successfully amplify.

To prepare segments for assembly, 3ul of each PCR reaction for segments 2-33 was pooled and purified using the Zymo Clean & Concentrator - 5 kit. The entirety of the pooled products were transformed for assembly. Amplicons 1 and 34 were fused by PCR assembly to sequences corresponding to the linearized ends of the assembly vector to create chimeric segments overlapping the vector. These reactions were purified and added to the assembly transformation seperately.

### Assembly transformation

The pooled assembly segments and assembly vector were transformed using a high efficiency yeast transformation method into the yeast strain BY4741. Briefly, cells are grown to mid-log phase before being collected and washed with lithium acetate. Cells are incubated further with PEG, lithium acetate, carrier DNA, assembly DNA, and DMSO, before being collected again and resuspended in calcium chloride before plating. Transformed cells were plated minimal selective media and allowed to grow at 30°C for ∼3 days before screening for succesful assembly.

### Identifying successfully assembly clones

Initial screening for assembly clones was performed using a high-throughput automated RT-PCR workflow described previously (30). Briefly, 96 colonies arising from the assembly transformation were selected and grown in liquid culture before isolating genomic DNA (also containing the assemblon YAC DNA). Isolated genomic DNA was used as template in microscale RT-PCR reactions on a Roche 1536 lightcycler, and monitored for amplification using the above described junction-spanning primers. PCR data from the lightcycler was used to calculate the cycle at which the fluoresence passed the threshold (the ‘crossing-point’ cycle). Clones having positive reactions for all assembly junctions were isolated and subjected to whole-genome sequencing of the yeast strain, mapping reads to a hybrid human and yeast reference genome.

## Acknowledgements

This work was supported in part by an NHGRI CEGS grant 1RM1HG009491. J.D.B is a founder and Director of the following: Neochromosome, Inc., the Center of Excellence for Engineering Biology, and CDI Labs, Inc. and serves on the Scientific Advisory Board of the following: Modern Meadow, Inc., Recombinetics, Inc., and Sample6, Inc.

## References

1. Friedman DS, O’Colmain BJ, Munoz B, Tomany SC, McCarty C, de Jong PT, et al. Prevalence of agerelated macular degeneration in the United States. Arch Ophthalmol. 2004;122(4):564–72.

2. Klein R, Klein BE, Linton KL. Prevalence of age-related maculopathy. The Beaver Dam Eye Study. Ophthalmology. 1992;99(6):933–43.

3. Vingerling JR, Dielemans I, Hofman A, Grobbee DE, Hijmering M, Kramer CF, et al. The prevalence of age-related maculopathy in the Rotterdam Study. Ophthalmology. 1995;102(2):205–10.

4. Rattner A, Nathans J. Macular degeneration: recent advances and therapeutic opportunities. Nat Rev Neurosci. 2006;7(11):860–72.

5. Bressler NM. Early detection and treatment of neovascular age-related macular degeneration. J Am Board Fam Pract. 2002;15(2):142–52.

6. Clemons TE, Milton RC, Klein R, Seddon JM, Ferris FL, 3rd, Age-Related Eye Disease Study Research G. Risk factors for the incidence of Advanced Age-Related Macular Degeneration in the Age-Related Eye Disease Study (AREDS) AREDS report no. 19. Ophthalmology. 2005;112(4):533–9.

7. Brown DM, Kaiser PK, Michels M, Soubrane G, Heier JS, Kim RY, et al. Ranibizumab versus verteporfin for neovascular age-related macular degeneration. N Engl J Med. 2006;355(14):1432–44.

8. Rosenfeld PJ, Brown DM, Heier JS, Boyer DS, Kaiser PK, Chung CY, et al. Ranibizumab for neovascular age-related macular degeneration. N Engl J Med. 2006;355(14):1419–31.

9. Smith W, Assink J, Klein R, Mitchell P, Klaver CC, Klein BE, et al. Risk factors for age-related macular degeneration: Pooled findings from three continents. Ophthalmology. 2001;108(4):697–704.

10. Edwards AO, Ritter R, 3rd, Abel KJ, Manning A, Panhuysen C, Farrer LA. Complement factor H polymorphism and age-related macular degeneration. Science. 2005;308(5720):421–4.

11. Haines JL, Hauser MA, Schmidt S, Scott WK, Olson LM, Gallins P, et al. Complement factor H variant increases the risk of age-related macular degeneration. Science. 2005;308(5720):419–21.

12. Fagerness JA, Maller JB, Neale BM, Reynolds RC, Daly MJ, Seddon JM. Variation near complement factor I is associated with risk of advanced AMD. Eur J Hum Genet. 2009;17(1):100–4.

13. Gold B, Merriam JE, Zernant J, Hancox LS, Taiber AJ, Gehrs K, et al. Variation in factor B (BF) and complement component 2 (C2) genes is associated with age-related macular degeneration. Nat Genet. 2006;38(4):458–62.

14. Maller JB, Fagerness JA, Reynolds RC, Neale BM, Daly MJ, Seddon JM. Variation in complement factor 3 is associated with risk of age-related macular degeneration. Nat Genet. 2007;39(10):1200–1.

15. Weeks DE, Conley YP, Tsai HJ, Mah TS, Schmidt S, Postel EA, et al. Age-related maculopathy: a genomewide scan with continued evidence of susceptibility loci within the 1q31, 10q26, and 17q25 regions. Am J Hum Genet. 2004;75(2):174–89.

16. Rivera A, Fisher SA, Fritsche LG, Keilhauer CN, Lichtner P, Meitinger T, et al. Hypothetical LOC387715 is a second major susceptibility gene for age-related macular degeneration, contributing independently of complement factor H to disease risk. Hum Mol Genet. 2005;14(21):3227–36.

17. Yang Z, Camp NJ, Sun H, Tong Z, Gibbs D, Cameron DJ, et al. A variant of the HTRA1 gene increases susceptibility to age-related macular degeneration. Science. 2006;314(5801):992–3.

18. Dewan A, Liu M, Hartman S, Zhang SS, Liu DT, Zhao C, et al. HTRA1 promoter polymorphism in wet age-related macular degeneration. Science. 2006;314(5801):989–92.

19. Yang Z, Tong Z, Chen Y, Zeng J, Lu F, Sun X, et al. Genetic and functional dissection of HTRA1 and LOC387715 in age-related macular degeneration. PLoS Genet. 2010;6(2):e1000836.

20. Kanda A, Chen W, Othman M, Branham KE, Brooks M, Khanna R, et al. A variant of mitochondrial protein LOC387715/ARMS2, not HTRA1, is strongly associated with age-related macular degeneration. Proc Natl Acad Sci U S A. 2007;104(41):16227–32.

21. Fritsche LG, Loenhardt T, Janssen A, Fisher SA, Rivera A, Keilhauer CN, et al. Age-related macular degeneration is associated with an unstable ARMS2 (LOC387715) mRNA. Nat Genet. 2008;40(7):892–6.

22. Dymond JS, Richardson SM, Coombes CE, Babatz T, Muller H, Annaluru N, et al. Synthetic chromosome arms function in yeast and generate phenotypic diversity by design. Nature. 2011;477(7365):471–6.

23. Annaluru N, Muller H, Mitchell LA, Ramalingam S, Stracquadanio G, Richardson SM, et al. Total synthesis of a functional designer eukaryotic chromosome. Science. 2014;344(6179):55–8.

24. Mitchell LA, Wang A, Stracquadanio G, Kuang Z, Wang X, Yang K, et al. Synthesis, debugging, and effects of synthetic chromosome consolidation: synVI and beyond. Science. 2017;355(6329).

25. Richardson SM, Mitchell LA, Stracquadanio G, Yang K, Dymond JS, DiCarlo JE, et al. Design of a synthetic yeast genome. Science. 2017;355(6329):1040–4.

26. Shen Y, Wang Y, Chen T, Gao F, Gong J, Abramczyk D, et al. Deep functional analysis of synII, a 770- kilobase synthetic yeast chromosome. Science. 2017;355(6329).

27. Wu Y, Li BZ, Zhao M, Mitchell LA, Xie ZX, Lin QH, et al. Bug mapping and fitness testing of chemically synthesized chromosome X. Science. 2017;355(6329).

28. Xie ZX, Li BZ, Mitchell LA, Wu Y, Qi X, Jin Z, et al. “Perfect” designer chromosome V and behavior of a ring derivative. Science. 2017;355(6329).

29. Zhang W, Zhao G, Luo Z, Lin Y, Wang L, Guo Y, et al. Engineering the ribosomal DNA in a megabase synthetic chromosome. Science. 2017;355(6329).

30. Mitchell LA, McCulloch LH, Pinglay S, Berger H, Bulajic M, Martin JA, et al. De novo assembly, delivery and expression of a 101 kb human gene in mouse cells. 2018.

31. Friedrich U, Myers CA, Fritsche LG, Milenkovich A, Wolf A, Corbo JC, et al. Risk- and non-risk-associated variants at the 10q26 AMD locus influence ARMS2 mRNA expression but exclude pathogenic effects due to protein deficiency. Human Molecular Genetics. 2011;20(7):1387–99.

32. Jones A, Kumar S, Zhang N, Tong Z, Yang J-H, Watt C, et al. Increased expression of multifunctional serine protease, HTRA1, in retinal pigment epithelium induces polypoidal choroidal vasculopathy in mice. Proceedings Of The National Academy Of Sciences Of The United States Of America. 2011;108(35):14578–83.

33. MacArthur J, Bowler E, Cerezo M, Gil L, Hall P, Hastings E, et al. The new NHGRI-EBI Catalog of published genome-wide association studies (GWAS Catalog). Nucleic Acids Research. 2017;45(D1):D896–D901.

34. Maurano MT, Humbert R, Rynes E, Thurman RE, Haugen E, Wang H, et al. Systematic localization of common disease-associated variation in regulatory DNA. Science. 2012;337(6099):1190–5.

35. Wallace HA, Marques-Kranc F, Richardson M, Luna-Crespo F, Sharpe JA, Hughes J, et al. Manipulating the mouse genome to engineer precise functional syntenic replacements with human sequence. Cell. 2007;128(1):197–209.

36. Wilson MD, Barbosa-Morais NL, Schmidt D, Conboy CM, Vanes L, Tybulewicz VL, et al. Species-specific transcription in mice carrying human chromosome 21. Science. 2008;322(5900):434–8.

